# ScopeViewer: A Browser-Based Solution for Visualizing Spatial Transcriptomics Data

**DOI:** 10.1101/2023.07.24.549256

**Authors:** Danni Luo, Sophie Robertson, Yuanchun Zhan, Ruichen Rong, Shidan Wang, Xi Jiang, Sen Yang, Suzette Palmer, Liwei Jia, Qiwei Li, Guanghua Xiao, Xiaowei Zhan

## Abstract

**Motivation:** Spatial transcriptomics (ST) enables a high-resolution interrogation of molecular characteristics within specific spatial contexts and tissue morphology. Despite its potential, visualization of ST data is a challenging task due to the complexities in handling, sharing and visualizing large image datasets together with molecular information.

**Results:** We introduce ScopeViewer, a browser-based software designed to overcome these challenges. ScopeViewer offers the following functionalities: (1) It visualizes large image data and associated annotations at various zoom levels, allowing for intricate exploration of the data; (2) It enables dual interactive viewing of the original images along with their annotations, providing a comprehensive understanding of the context; (3) It displays spatial molecular features with optimized bandwidth, ensuring a smooth user experience; and (4) It bolsters data security by circumventing data transfers.

**Availability:** ScopeViewer is available at: https://datacommons.swmed.edu/scopeviewer

**Supplementary information:** Supplementary data are available at Bioinformatics online.

## 1. Introduction

Spatial transcriptomics (ST) technologies have made significant advancements in recent years (Zhang, et al., 2021). ST techniques offer high-resolution transcriptome measurements with spatial information within tissues, thereby opening new avenues for understanding cellular and molecular spatial distributions (Crosetto, et al., 2015), and their associated links to diseases (Shah, et al., 2018). A typical ST dataset pairs with high-resolution images, often comprising millions of pixels. This facilitates a dual visualization of cellular and tissue structures alongside quantitative molecular features, including gene expression and protein abundance. Examining molecular characteristics within spatial and morphological contexts could pave the way for new biological discoveries. A comprehensive tool for visualizing ST data will streamline data exploration and analysis, aiding researchers in comprehending molecular features within specific biological contexts.

Working with high-resolution tissue images and ST data introduces significant challenges due to the huge volume of these datasets. Consider a standard pathology image of 20,000 by 20,000 pixels, which can amass a file size of approximately one gigabyte. This large size complicates both the image’s transfer and visualization, often requiring specialized software tools. Moreover, there is a high degree of complexity inherent in visualizing high-dimensional molecular features alongside the intricate cell and tissue structures. As a result, current software packages often struggle to simultaneously display molecular details with corresponding high-resolution tissue images effectively. Further complications arise from software interfaces that require tedious manual input from users to toggle visibility of data layers. A more streamlined solution might include offering a dual-view approach. This would allow users to see the data layer in one view, while simultaneously hiding it in another, with synchronized panning and zooming capabilities. Lastly, the ability for researchers to explore ST data on their own systems, without uploading or sharing data externally, is an important consideration. This not only bolsters data security but also enhances user accessibility. To address these prevalent challenges, we developed ScopeViewer, a browser-based visualization software, available at: https://datacommons.swmed.edu/scopeviewer.

## 2. Methods and Results

ScopeViewer operaters as a web application, requiring nothing more than a web browser for its execution. It was designed using ReactJS JavaScript framework. To utilize ScopeViewer, users simply navigate to the website and input the image information and ST data from a local path, using the JSON syntax. ScopeViewer generates an interactive user interface directly within the web browser. The platform’s design leverages the versatility of web browsers, thereby eliminating the need for users to install specific software on their hardware.

### 2.1 Supports for multiple annotation formats

When conducting pathology image analysis or exploring spatial transcriptomics data, it is crucial to view the image at varying magnification levels. Additionally, users often need to overlay various annotations. These might include (1) tissues from disparate anatomical locations; and (2) spatial spots generated by the 10x platform. To accommodate these needs, ScopeViewer incorporates the widely used deep zoom format (a Microsoft-maintained XML specification for viewing large images) and the advanced OpenSeaDragon platform (OpenSeaDragon, 2023). ScopeViewer’s functionality extends beyond displaying multiple layers of large images at different magnification levels. It also supports dual views, a feature that enables side-by-side synchronized display. Additionally, ScopeViewer accommodates user annotations of various shapes, including lines, rectangles, ellipses, and polygons. Users can conveniently specify these layers within an online JSON editor, which provides instant feedback for any syntax errors. This feature enhances user-friendliness and ensures accurate data input.

### 2.2 Reduction of data transfer using a transpiled SQLite module

Spatial transcriptomics (ST) generates both high-resolution tissue image data and high-dimensional spatial molecular data, resulting in large datasets that are difficult to browse over the Internet. The process of transferring and processing such extensive ST data can be time-consuming and challenging. To overcome this obstacle, we incorporated a tailored SQLite database implementation that offers two key features: (1)a WebAssembly version of SQLite. This was transpiled from its original C codes and provides high execution speed within the browser, which significantly enhances performance (Andrés and Pérez, 2017; Rossberg, 2021). (2) It has the capacity to fetch expression quantities from the SQLite database through HTTP Byte-Range headers. This functionality minimizes data transfer from the SQLite Virtual File System (VFS), making it more efficient. As a result of these optimizations, the webpage size is reduced from 180M (original molecular data) to 17.2M without cache, or 6.5M with the cache. This inventive approach simplifies the visualization of spatial transcriptomics data within browser-based applications, enhancing user experience and enabling more effective research analysis.

### 2.3 Application: a breast cancer data from the 10X Visium platform

We demonstrate the use of ScopeViewer through a 10X Visium breast cancer FFPE sample (Janesick, et al., 2022). To analyze the cell types and spatial distribution, we applied HD-Yolo, a deep-learning cell segmentation and classification algorithm on the whole slide image (Rong, et al., 2023). The image file measures 25,233 x 27,452 pixels and is 143M in size. Both the original image and its annotations are stored in DZI formats and deployed as the default ScopeViewer Demo to showcase three key features: (1) The original H&E slides are displayed on the left, while the algorithm-annotated image appears on the right within a synchronized interface (Fig. 1A). At the highest magnification level, tumor nuclei, necrosis, red blood cells, and stroma cells are annotated in green, cyan, magenta, and red, respectively. (2) The molecular transcriptome data (the spots) can be overlaid on the image (Fig. 1B). This shows that the cancer biomarker gene FASN is expressed highly in the tumor region, clustering with tumor cells (He, et al., 2020). (3) No genomic data are transferred to the ScopeViewer web server, as data exchanges occur solely between the browser and the data server (Fig. 1C). This means the ScopeViewer web server instructs the user’s browser to retrieve and display relevant information, without accessing potentially sensitive data, as its website does not communicate with the data server.

**Figure 1:**
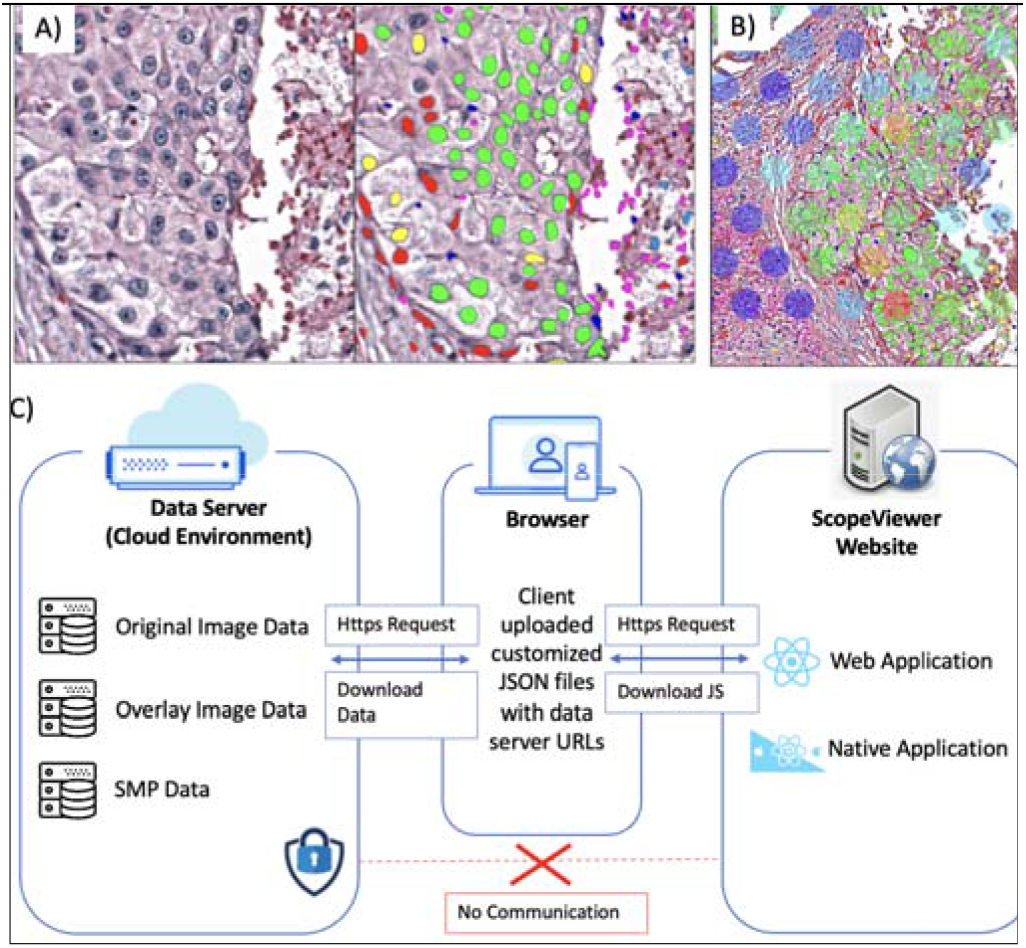
ScopeViewer for visualization ST data. (1) synchronized views for H&E pathology image and AI facilitated cell segmentat (2) efficiently overlaying spatial transcriptomics features (here sh FASN, a breast cancer biomarker gene); (3) visualization will not genomic data to the ScopeViewer webserver.

## 3. Conclusion

ScopeViewer is feature-rich, cloud-based, and secure tool for visualizing ST datasets. We envision that it will be a valuable resource **for** data exploration and sharing within the wider research community.

## Funding

This work has been supported by the National Institutes of Health [R01GM140012, R01GM141519, R01DE030656, U01CA249245, U01AI169298, R01GM126479], the Cancer Prevention and Research Institute of Texas [CPRIT RP180805 and RP230330] and National Science Foundation [2113674, 2210912].

## Conflict of Interest

none declared.

## Notes

https://datacommons.swmed.edu/scopeviewer

### Competing Interest Statement

The authors have declared no competing interest.

