## Supplementary Materials for "ScopeViewer: A Browser-Based Solution for Visualizing Spatial Transcriptomics Data"

**Supplementary Texts**

1. **A common file format that supports multiple spatial transcriptomics platforms**

**An overview of spatial transcriptomics platforms**

Cellular and molecular spatial organizations play essential roles in biological functions. The development of spatial transcriptomics (ST) techniques have achieved significant breakthroughs in recent years (Zhang, et al., 2021). These techniques enable high resolution transcriptome measurement with tissue spatial information, which provides new opportunities to advance our understanding of cellular and molecular spatial distributions (Crosetto, et al., 2015) and their relationships with diseases (Shah, et al., 2018). Popular ST techniques can be classified into two types: imaging-based and sequencing-based. Based on the single-molecule fluorescence *in situ* hybridization (FISH), the two most classical imaging-based ST techniques are sequential FISH (seqFISH) (Lubeck, et al., 2014) and multiplexed error-robust FISH (MERFISH) (Chen, et al., 2015). These techniques can measure the expression levels of hundreds to thousands of genes in individual cells. STARmap is a recently developed 3D intact-tissue RNA sequencing approach with single-cell resolution (Wang, et al., 2018). Sequencing-based techniques, such as spatial transcriptomics (Stahl, et al., 2016), 10x Visium platform, and high-definition spatial transcriptomics (HDST) (Vickovic, et al., 2019) use spatial barcode probes to capture RNA molecules and then synthesize and sequence their complementary DNA molecules. These techniques quantify the expression levels of the entire transcriptome by utilizing hundreds to thousands of barcode probes, referred to as spots, which collectively capture the profiles of cell. In summary, all the above technologies can quantify transcriptomics in a spatial context and we found that it is possible to store the measurements in a unified format for fast web-based visualizations.

**Data preparation**

ScopeViewer has the ability to generate spatial molecular profiling (SMP) layer, which overlays spot data on the histology image. To achieve this function, users need to provide corresponding SMP data for the image, which includes three essential files: 1) a gene expression matrix $Y$ with $P\times N$ dimensions, where $N$ denotes the number of spots or cells and $P$ denotes the number of genes. Each entry $y_{ij}$ is the read count for gene $i$ collected at spot/cell $j$; 2) a location table $T$ with $N$ rows and two columns. Each row contains the x and y coordinate of each spot/cell; and 3) a vector with length $P$ containing the list of gene names. All the data tables above need to be stored in “.csv” format. Users need to use the provided Python script to convert all files into three tables (‘smp_count_sql’, ‘smp_loc_sql’, and ‘gene_list_sql’) in a SQlite database file, and then provide the path of generated .db file with "smp_layer" key in a .json file. Detailed instructions and example files can be found on our website.

2. **Transpile SQLite as a web module**

Spatial transcriptomics datasets often range from megabytes to gigabytes in size. While this scale of disk space is easily manageable for desktop-based visualization software, it poses a significant challenge for cloud-based applications like ScopeViewer due to the slow transfer of large data volumes from the internet to the user's browser.

To overcome this obstacle, we adopted the use of the SQLite database, transpiled to WebAssembly (WASM) for efficient binary execution and enhanced computational speed. We further leveraged an open-source implementation to selectively retrieve B-tree blocks from the underlying SQLite database file, thereby transferring only necessary blocks using the byte range feature commonly found in web servers or cloud storage services (e.g., Amazon S3).

This process results in transferring only a small fraction of transcriptomics data visible to the user over the internet. In our benchmarks, we typically observed a reduction in data transfer from 1/10th to 1/100th of the original data, demonstrating that our transpiled SQLite approach substantially decreases the bandwidth required for visualizing spatial transcriptomics datasets.

**Supplementary Figure 1. ScopeViewer provides a convenient JSON editor for customized visualization.** The ScopeViewer tool enables customizable visualization of spatial transcriptomics data via a JSON configuration file. We provide an editing feature supported by a JavaScript implementation based on the Monaco editor (under the MIT license), showcasing the following capabilities: (1) Syntax colorization; (2) Real-time syntax validation; (3) Collapsible code blocks; (4) Bracket matching for improved readability and error minimization.

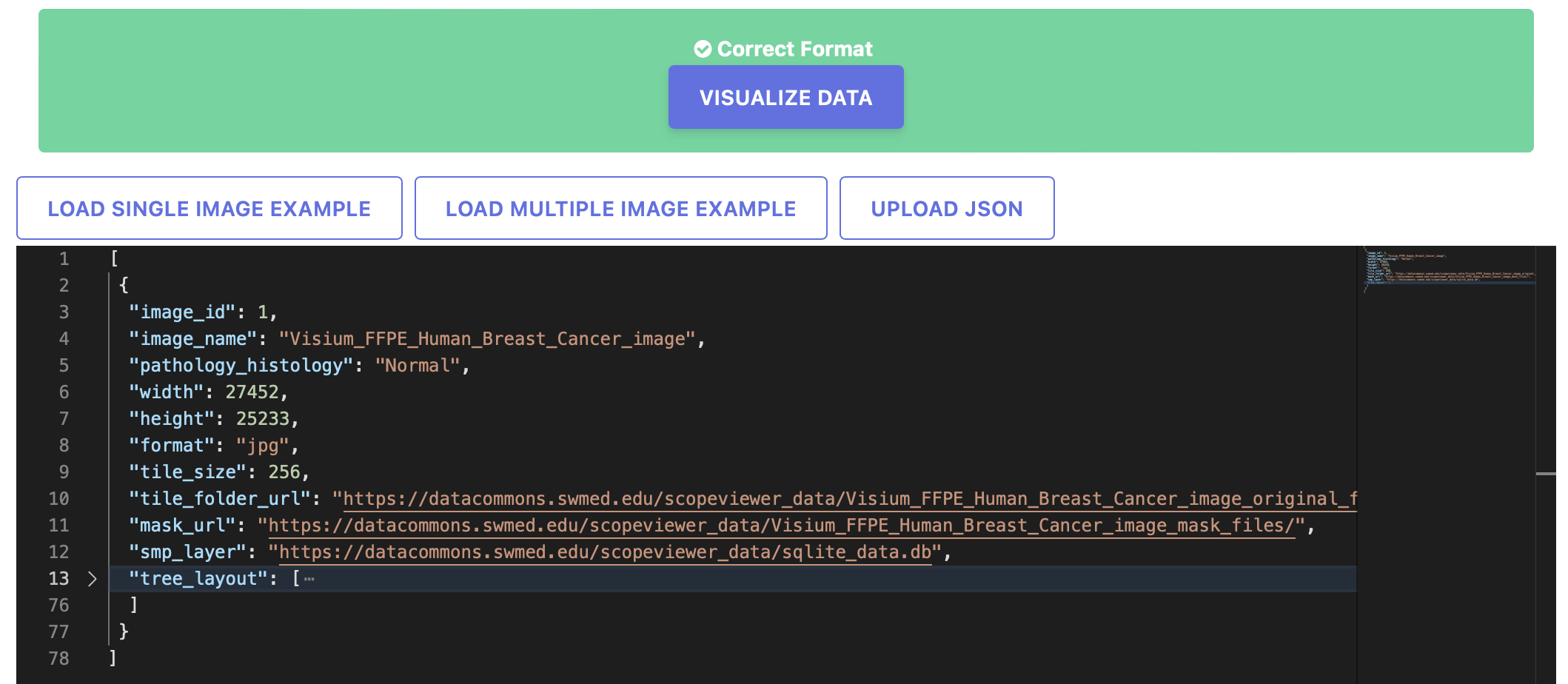

**Supplementary Figure 2. ScopeViewer allows visualization of annotations in a**

**hierarchical structure.** In tissue slides, multiple types of cells exist, and they can have a multiple level hierarchical structure (e.g., cell lineages). ScopeViewer allow users to specify such tree-structure in the JSON file. This illustration presents five distinct annotation layers (left) derived from the corresponding JSON source codes (right). Users have the flexibility to selectively display or conceal individual layers as required.

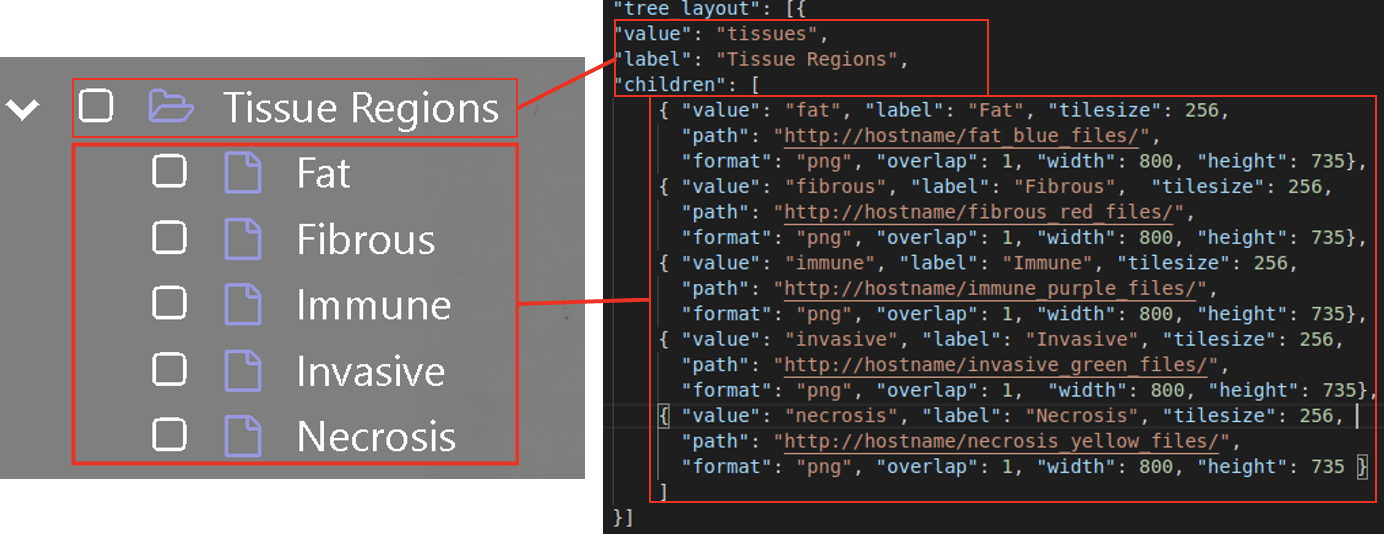
